# Genome-scale model of *Rhodotorula toruloides* metabolism

**DOI:** 10.1101/528489

**Authors:** Ievgeniia A Tiukova, Sylvain Prigent, Jens Nielsen, Mats Sandgren, Eduard J Kerkhoven

**Affiliations:** Systems and Synthetic Biology, Department of Biology and Biological Engineering, Chalmers University of Technology, Gothenburg, Sweden; Department of Molecular Sciences, Swedish University of Agricultural Sciences, Uppsala, Sweden; UMR 1332 BFP, Univ Bordeaux, F33883 Villenave d’Ornon, France

**Keywords:** genome-scale model, yeast, *Rhodotorula toruloides*, metabolism

## Abstract

The basidiomycete red yeast *Rhodotorula toruloides* is a promising platform organism for production of biooils. We present *rhto-GEM*, the first genome-scale model (GEM) of *R. toruloides* metabolism, that was largely reconstructed using RAVEN toolbox. The model includes 852 genes, 2731 reactions, and 2277 metabolites, while lipid metabolism is described using the SLIMEr formalism allowing direct integration of lipid class and acyl chain experimental distribution data. The simulation results confirmed that the *R. toruloides* model provides valid growth predictions on glucose, xylose and glycerol, while prediction of genetic engineering targets to increase production of linolenic acid, triacylglycerols and carotenoids identified genes – some of which have previously been engineered to successfully increase production. This renders *rtho-GEM* valuable for future studies to improve the production of other oleochemicals of industrial relevance including value-added fatty acids and carotenoids, in addition to facilitate system-wide omics-data analysis in *R. toruloides*. Expanding the portfolio of GEMs for lipid accumulating fungi contributes to both understanding of metabolic mechanisms of the oleaginous phenotype but also uncover particularities of the lipid production machinery in *R. toruloides*.

## 1 Introduction

*Rhodotorula toruloides* (syn. *Rhodosporidium toruloides*) is a basidiomycetous yeast belonging to the subphylum Pucciniomycotina and occurs naturally in a wide range of habitats including surfaces of leaves, soil and sea water (Sampaio, 2011). The broad substrate range of *R. toruloides* and its ability to accumulate lipids exceeding half of its cell dry weight has made this yeast a popular system for production of biological oils from inedible substrates (e.g. pentose sugars, crude glycerol) (Park, Nicaud, & Ledesma-Amaro, 2018). The ready scalability and fast rate of lipid production by oleaginous yeasts such as *R. toruloides* makes this approach a competitive alternative to emerging “third generation” biodiesel produced from carbon dioxide. A commercially viable process of biodiesel production using *R. toruloides* was recently patented in Brazil where conversion of a mixture of sugarcane juice and urea into lipids achieved a productivity of 0.44 g/L h (Soccol et al., 2017). The *R. toruloides* lipid fraction contains ω-3 linolenic acid and heptadecenoic acid which makes this yeast a promising organism for the production of pharma- and nutraceuticals at the same time as it is also a natural producer of several carotenoid pigments including torularhodin, torulene, γ-carotene and β-carotene (Buzzini et al., 2007).

One of the major determinants of the oleaginous phenotype of *R. toruloides* is its capacity for acetyl-CoA production (Zhu et al., 2012). Unlike non-oleaginous yeasts such as the baker’s yeast *Saccharomyces cerevisiae, R. toruloides* possesses the enzyme ATP:citrate lyase, which is the main source of acetyl-CoA for lipid synthesis. In addition, a mitochondrial beta-oxidation pathway provides additional source of acetyl-CoA in this yeast (Vorapreeda, Thammarongtham, Cheevadhanarak, & Laoteng, 2012). Lipid biosynthetic reactions downstream of acetyl-CoA synthesis do not differ between oleaginous and non-oleaginous yeast species, simplifying the use template models from non-oleaginous yeasts.

A better understanding of lipid metabolism in *R. toruloides* has applications beyond the production of lipids for fuel and nutraceuticals. *R. toruloides* has also been used in screens for novel anti-obesity drugs (Lee, Cheon, & Rhee, 2018), while lipid synthesis has been shown to be a promising target for antifungal therapies (Pan, Hu, & Yu, 2018). Several species within the *Rhodotorula* genus have been identified as emerging pathogens in both animals and people (Wirth & Goldani, 2012). Intracellular glycerol accumulation caused by hydrolysis of storage triglycerides by pathogenic fungi has been shown to play a crucial role in turgor generation for penetration and invasion of tissues (Nguyen et al., 2011; C. Wang & St. Leger, 2007). Modelling of *R. toruloides* lipid metabolism can therefore also aid in unravelling the pathobiology of this group of yeasts.

Genome-scale metabolic models (GEMs) are comprehensive summaries of the metabolic network in an organism, which are derived from the available genome sequence. GEMs can be used for identification of potential metabolic engineering targets, or select the best performing metabolic network among alternatives (Kerkhoven, Lahtvee, & Nielsen, 2015). Extending the portfolio of GEMs for various organisms can enhance the benefits of bioprospecting and aid in the design and improvement of performance of synthetic organisms. The generation of a genome scale metabolic model of *R. toruloides* lipid synthesis would provide a deeper understanding of oleaginous ability and facilitate genetic engineering strategies. Although a number of metabolic models of lipid production in *R. toruloides* have been reported to date (Bommareddy, Sabra, Maheshwari, & Zeng, 2015; Castañeda, Nuñez, Garelli, Voget, & De Battista, 2018), this study represents the first genome-scale model of its metabolism.

## 2 Materials and Methods

### 2.1 Draft model reconstruction

The genome-scale model, named *rhto-GEM*, is based on the genome sequence of *R. toruloides* strain NP11 (Zhu et al., 2012). The reconstruction of *rhto-GEM* was primarily performed using RAVEN 2.2.1, a MATLAB toolbox for genome-scale model reconstruction (H. Wang et al., 2018). All steps of the reconstruction are documented in detail on the GitHub repository under the folder *ComplementaryScripts/reconstruction*. More information on the GitHub repository is provided below.

To reconstruct those parts of metabolism that are relatively conserved between fungal species, the well-curated GEMs of *Saccharomyces cerevisiae* (yeast-GEM version 8.2.0, doi:10.5281/zenodo.1495483, (Lu et al., 2019)) and *Yarrowia lipolytica* (iYali4.1.1, (Kerkhoven, Pomraning, Baker, & Nielsen, 2016)) were taken as template models, while orthologous genes were identified via bi-directional BLASTP (Camacho et al., 2009) against the *S. cerevisiae* S288c and *Y. lipolytica* CLIB 122 reference genomes. All 1-1 orthologs were included, after cut-offs of E-value <1e-20; identity >35% and alignment length >150 bp. Additional orthologs between *R. toruloides* and *S. cerevisiae* were identified as provided by MetaPhOrs (Pryszcz, Huerta-Cepas, & Gabaldón, 2011), filtered for ortholog pairs with confidence scores of 1 and whose PhylomeDB tree contained at least two members.

### 2.2 Gap-filling with Meneco

To obtain a functional model, a gap-filling step was performed to add reactions necessary to produce biomass from the preferred growth medium of *R. toruloides*. To avoid self-producing loops due to stoichiometric inconsistencies, we utilized Meneco 1.5.2 (Prigent et al., 2017) in combination with yeast-GEM (v. 8.2.0) as database of repair reactions. In Meneco, target compounds correspond to metabolites present in the biomass function, while seed compounds are composed of metabolites present in the growth medium, plus some cofactors and metabolites required for FBA growth. If no path exists between seed and target compounds, Meneco proposes one minimal set of reactions (or several minimal sets of same size) coming from a database of reactions to fill the gaps. The sets of seed and target compounds are given in on the GitHub repository under the folder *ComplementaryScripts/meneco*. The seed compounds included uncharged tRNAs as the biomass reaction explicitly represents protein translation as the transfer of amino acids from tRNAs. The union of all proposed completions was included in the draft model, while manual curation was performed to confirm the likelihood of those reactions and to identify their corresponding genes in *R. toruloides*.

### 2.3 Lipid metabolism with SLIMEr

Lipid metabolism was described using the SLIMEr formalism, which Splits Lipids Into Measurable Entities (Sánchez, Li, Kerkhoven, & Nielsen, 2019). To minimize the number of unique lipid species, we inferred from experimental data (Wei, Siewers, & Nielsen, 2017) which acyl chains can be expected at each position: in phospholipids and TAGS the sn-1 position is populated by saturated acyl chains (i.e. 16:0 or 18:0); sn-2 positions by unsaturated acyl chains (i.e. 18:1, 18:2 or 18:3) and sn-3 positions by saturated or monounsaturated acyl chains (i.e. 16:0, 18:0, 18:1). Palmitoleate (16:1) is not modelled as it is only a minor contributor (< 5%) to the overall acyl chain distribution (Tiukova et al., 2019). Cardiolipin maturation is furthermore simplified by assuming that the monolysocardiolipin acyltransferase only utilizes phosphatidylcholine with acyl configuration 1-16:0; 2-18:1 as co-substrate. Consequently, this resulted in 67 SLIME reactions and 1022 curated reactions in lipid metabolism, instead of 920 and 7130 reactions if all combinations of the five acyl chains (i.e. 16:0; 18:0; 18:1; 18:2; 18:3) were allowed. To facilitate adjusting the lipid composition in the model by considering measured lipid class and acyl chain distributions, we provide the functions *adjustRhtoBiomass* and *scaleLipidsRhto* in the *ComplementaryScripts/experimental* folder.

### 2.4 Further model development and distribution via GitHub repository

The growth-associated and non-growth associated energy requirements were fit to measured glucose uptake rates from continuous cultivations of *R. toruloides* as reported in literature (Shen et al., 2013), and set at 132.7 mmol gDCW^−1^ and 3.39 mmol (gDCW h)^−1^, respectively. The biomass composition was modified from yeast-GEM to include *R. toruloides* lipid class and acyl chain distributions, as provided in *ComplementaryData/data*. The *consumeSomething* and *produceSomething* functions from RAVEN were used to ensure there is no net production or consumption of mass by any reaction in the model. The *rhto-GEM* model is hosted on a dedicated GitHub repository (http://github.com/SysBioChalmers/rhto-GEM). Here, all scripts for model reconstruction are provided, in addition to the model in various file formats, e.g. SBML, YAML, TXT and XLSX, and scripts for performing the simulations detailed in this manuscript. This environment allows for versioning and comparison of the model, reporting and tracking of issues, organization of development and continuous integration. Memote 0.9.2. is a model test suite (Lieven et al., 2018) used to assess model quality, which is automatically run via Travis CI with each new model release, currently skipping the consistency tests due to their long duration.

### 2.5 Model simulations

Flux balance analysis was performed with RAVEN toolbox, using constraints on exchange fluxes as specified in the text, while also detailed in the relevant scripts in the *ComplementaryScripts* folder. All simulations here were performed with *rhto-GEM* version 1.2.1. Gene essentiality was predicted using *singleGeneDeletion* from COBRA toolbox 3.0.6 (Heirendt et al., 2017), where growth rates reduced by more than two-thirds were classified as lethal. Reactions significantly affecting TAG biosynthesis were identified using *singleRxnDeletion* from COBRA toolbox. To predict genetic targets for metabolic engineering the FSEOF (flux scanning of enforced objective function, (Choi, Lee, Kim, & Woo, 2010)) implementation of RAVEN was used. Exchange reactions were added for the products of interest, which were optimized with either glucose or xylose as carbon source. We performed FSEOF analysis using triacylglyceride (1-18:0, 2-18:1, 3-18:0) as a representative species for TAG, while this TAG is furthermore of interest as major component of cocoa butter. The slope parameter derived from FSEOF is indicative of how strong each reaction is contributing towards a shift from growth towards production of the target compound, and thereby suggests which reactions are promising targets for overexpression to increase productivity.

## 3 Results and Discussion

### 3.1 Step-wise reconstruction of rhto-GEM

To support the development of *R. toruloides* as promising microbial biocatalyst for oleochemical production, we developed a genome-scale model for *R. toruloides* strain NP11 through a combination of semi-automated reconstruction and manual curation based on literature data (**Fig 1A**). The reconstruction and curation process is tracked in a publicly accessible Git repository, an environment that allows for open, reproducible and trackable development and curation of genome-scale models by any member of the research community. Users are encouraged to report issues with the existing model and contribute to the continuous development, while the step-wise reconstruction described here is fully documented in the repository.

**Fig. 1.**
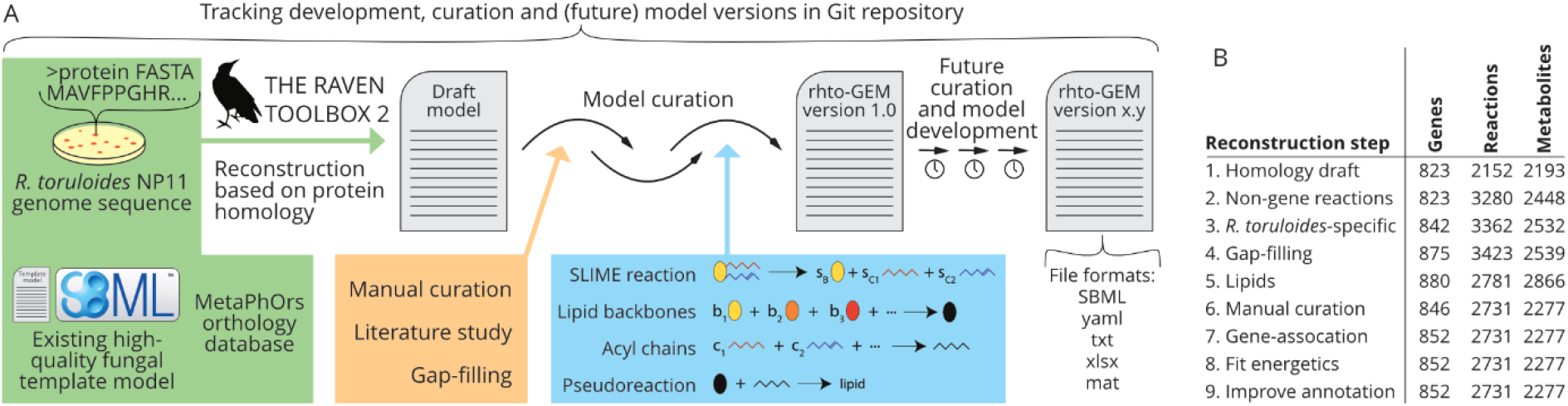
Development and distribution of *rhto-GEM*. (A) Schematic overview of the reconstruction pipeline, starting from generating a draft model using RAVEN, a protein FASTA of the *R. toruloides* genome and a template model. Through manual curation and inclusion of the SLIMEr formalism for lipid metabolism a functional model *rhto-GEM* is constructed, which is available in several different file formats. The whole development is tracked in a Git repository, which furthermore makes the model amenable to trackable future curation and development. (B) Overview of model metrics during the different steps of model development.

Bidirectional protein-blast (Camacho et al., 2009) with RAVEN (H. Wang et al., 2018) Querying of the *R. toruloides* and *S. cerevisiae* genomes by either BLAST or the phylogeny-based MetaPhOrs identified respectively 628 and 571 pairs of orthologous genes that were annotated to the template yeast-GEM model. In addition, querying the phylogeny-based ortholog repository MetaPhOrs yielded 571 pairs of orthologs model-annotated genes. Complementary to *S. cerevisiae* as a template model, also the GEM of oleaginous yeast *Y. lipolytica* (Kerkhoven et al., 2016) was queried for orthologous genes, resulting in the identification 22 additional orthologous genes. Combined, this first step of model reconstruction rapidly identified 823 *R. toruloides* genes connected to 2152 reactions (**Fig 1B**), representing parts of metabolism that are well conserved between fungal species.

In the second step of model reconstruction, a large number of non-gene associated reactions, primarily pseudo-reactions, exchange and intracellular transport reactions, were transferred from the template model to the draft reconstruction. While such reactions are required to obtain a functional model, not all the template-derived reactions are required to support growth in the final *R. toruloides* model. Therefore, unconnected non-gene associated reactions were removed from the model at a late stage of the reconstruction.

In addition to automated template-based reconstruction, it is imperative to curate organism-specific reactions and pathways to obtain a representative and comprehensive model. In the third step of *rhto-GEM* reconstruction we included 7 reactions from the carotene and torulene biosynthetic pathways (Buzzini et al., 2007), while fatty-acid degradation through mitochondrial beta-oxidation introduced 67 further reactions (Zhu et al., 2012). Along with synthesis and degradation pathways of C18:2 and C18:3 fatty acids, the third step of model reconstruction increased the reaction and gene count to 3362 and 842, respectively.

### 3.2 Topological-based gap-filling

As the resulting draft model was unable to support the production of biomass, we performed gap-filling as the fourth step in the reconstruction. For this we utilized Meneco (Prigent et al., 2017), a gap-filling tool that aims to find a topological path in the bipartite graph of reactions and compounds between seed and target compounds. By utilizing yeast-GEM as database of reactions, Meneco proposed that 13 reactions should be added to the model to make it functional. A total of 24 different sets of 13 reactions were identified, the union of these having a size of 19 reactions. As no information was available to select a particular set of reactions among the 24 possibilities, we added all 19 reactions. If any of the newly-added reactions were catalysed by an enzyme (i.e. annotated with a gene in the model), then we assumed that also other reactions that are catalysed by the same enzyme should be included in the draft model. Through this approach we added three more reactions. Addition of all 22 reactions sufficed for the model to produce biomass. In addition to the topological-based gap-filling, manual curation throughout the development process identified a number of additional genes and reactions, resulting in an extended draft model with 3423 reactions and 875 genes.

### 3.3 Representation of lipid metabolism

As particular interest on *R. toruloides* is focused on its oleaginous nature, attention was paid to accurately depict lipid metabolism. Recently, we have developed the SLIMEr formalism for describing lipids in genome-scale models, which Splits Lipids Into Measurable Entities (Sánchez et al., 2019). This formalism represents the flexibility of lipid metabolism while allowing incorporation of measurements of lipid classes and acyl chain distributions, briefly explained in the Methods section, while a detailed analysis of the practical implications of this approach is provided in (Sánchez et al., 2019).

In the fifth step of reconstruction we applied the SLIMEr formalism as previously described for *S. cerevisiae*. As the acyl chain distribution of *R. toruloides* is different from *S. cerevisiae*, e.g. the presence of C18:2 and C18:3, this required extensive manual curation of the SLIME reactions, culminating in a lipid-curated draft model with 2781 reactions, which is less than before curation of lipid metabolism as non-relevant reactions (based on lipid acyl chain compositions) were discarded. We subsequently populated the model with FAME and lipid class data obtained from mid-exponential phase bioreactor cultivations of *R. toruloides* on glucose (Tiukova et al., 2019).

### 3.4 Quality control and validation of rhto-GEM

To transform this functional draft model to the first version of the *R. toruloides* GEM, additional manual curation was performed where in step 6 reactions not connected to the main network, as introduced early in the reconstruction process, were removed. In step 7, remaining template-derived genes were replaced by their *R. toruloides* orthologs where possible and otherwise deprecated, while in step 8 of the reconstruction the (non-)growth associated maintenance energy was fitted to experimentally determined growth and glucose uptake rates (Shen et al., 2013), and in step 9 the annotation of metabolites and reactions was improved. The resulting model, version 1.2.0, is the first curated genome-scale model of *R. toruloides* with a total of 2731 reactions, 2277 metabolites and 852 genes (**Fig 1B**). To track model quality, each time a new model version is released a memote (Lieven et al., 2018) snapshot report is generated, by running a standardized set of metabolic model tests focusing on e.g. annotations and consistency. A full snapshot report of *rhto-GEM* resulted in a memote test score of 62% (Supplementary Material S1). In particular, a low stoichiometric consistency is reported with a memote score of nearly 40%. This is at least partially an artefact of the SLIME reactions whose product stoichiometries are weight-normalized, which allows for direct integration of lipid measurements that are provided as grams per gram dry cell weight.

We structurally compared rhto-GEM with previously reported small-scale models of *R. toruloides* (Bommareddy et al., 2015; Castañeda et al., 2018). and identified a number of manually curated gene associations and reactions that were differently defined in the semi-automatically reconstructed rhto-GEM. These changes were used to curate the model, to yield version 1.2.1 that was used for further analysis. The maximum theoretical TAG production yields in the small-scale models were slightly lower than the yield predicted from rhto-GEM, which can be attributed to the absence of complex I of the oxidative phosphorylation resulting in lower energy yields (**Fig 2A**).

**Fig. 2.**
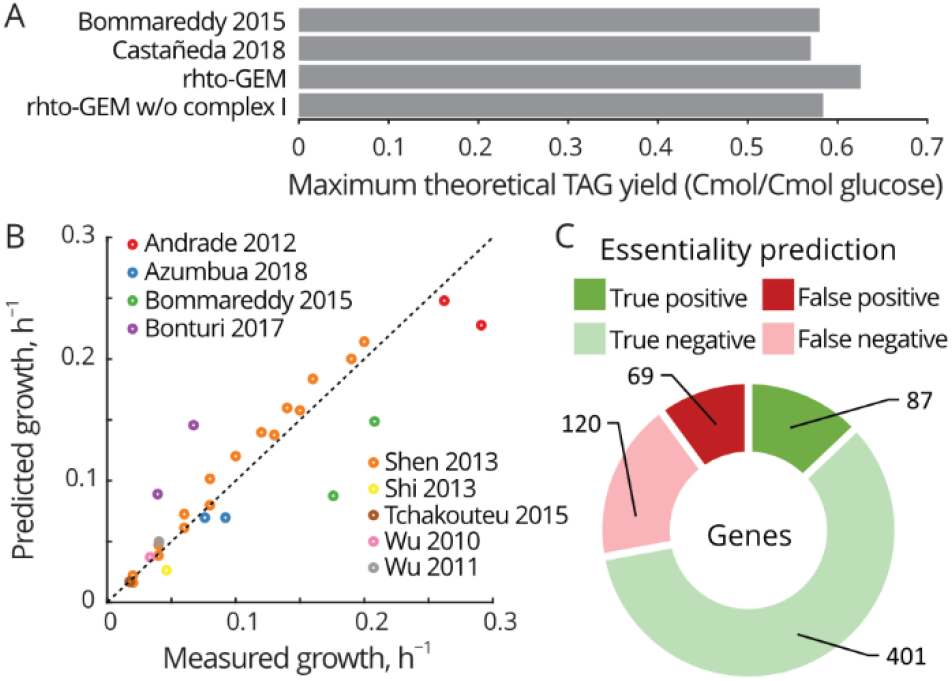
Growth and essentiality prediction from experimental data. (A) Maximum theoretical TAG yield on glucose as estimated with two small-scale models and *rhto-GEM* with and without complex I of the oxidative phosphorylation. (B) Comparison of experimentally measured growth rates and model predicted growth rates when constrained for the same carbon source uptake rates. Each data-point corresponds to one experimental growth measurement. The dotted line exemplifies perfect correlation. (C) Comparison of gene essentiality predictions from *rhto-GEM* with gene essentiality inferred from a T-DNA random insertion study (Coradetti et al., 2018). Shown are results from the 704 genes that are included in the model and that could be uniquely mapped to the T-DNA study.

To validate *rhto-GEM* functionality we compared its growth rate to experimental measured values. From literature we gathered glucose, glycerol and xylose uptake rates from *R. toruloides* bioreactor cultivations (Andrade, Leal, Roseiro, Reis, & da Silva, 2012; Azambuja, Bonturi, Miranda, & Gombert, 2018; Bommareddy et al., 2015; Bonturi, Crucello, Viana, & Miranda, 2017; Shen et al., 2013; Shi, Feng, Lee, & Chen, 2013; Tchakouteu et al., 2015; Wu, Hu, Jin, Zhao, & Zhao, 2010; Wu, Zhao, Shen, Wang, & Zhao, 2011), constrained the uptake rates in *rhto-GEM* to the reported values and optimized for biomass production. A good correlation could be found during growth on glucose, glycerol and xylose (**Fig 2B**). While the exact culture conditions, media compositions and *R. toruloides* strains used diverge between the literature-derived growth rates, these variables do not relate to how well they correlate with model-predicted growth rates. To evaluate the gene essentiality predictions, we used data from a recent study where a random mutant library was generated by T-DNA insertion in over 6,000 genes (Coradetti et al., 2018). A total 1,337 genes were identified as putative essential genes due to their recalcitrance to T-DNA insertion, of which 207 genes are associated to reactions in *rhto-GEM*. As it is unclear whether recalcitrance to T-DNA insertion is solely based on gene essentiality, we took the more conservative assumption that genes *not* recalcitrant to insertion are *not* essential. Simulations with *rhto-GEM* indicated a specificity (or true negative rate) of 0.853 (**Fig 2C**).

### 3.5 Essential reactions for oleaginous phenotype

*R. toruloides* is ranked as one of the best performing yeast species for lipid production together with the oleaginous yeasts *Lipomyces starkeyi* and *Trichosporon oleaginosus* (Papanikolaou & Aggelis, 2011). To evaluate which reactions are stoichiometrically most influential in the oleaginous phenotype of *R. toruloides*, we ran reaction essentiality analysis where the production of one particular triacylglycerol (TAG; a major storage lipid) was set as cellular objective and the resulting TAG yield was indicative of the essentiality of each reaction. Comparing oleaginous-essential reactions between three carbon sources (**Table 1**), the largest differences are related to the carbon assimilation pathway: pentose phosphate pathway reactions are more affecting for lipid production on xylose compared to glycerol. Acetyl-CoA carboxylase, which has previously been identified as an important target for increased TAG production (S. Zhang et al., 2016), was here identified as essential on all tested carbon sources. Many reactions that might be *perceived* to be essential were not identified as such in our analysis, such as the last step of TAG biosynthesis using a fatty acyl-CoA and diacylglycerol as catalysed by diacylglycerol acyltransferase. However, TAGs can alternatively be generated by reshuffling acyl chains between phospholipids and diacylglycerols. The relatively small number of essential reactions therefore demonstrates the high flexibility of lipid metabolism.

**Table 1.**
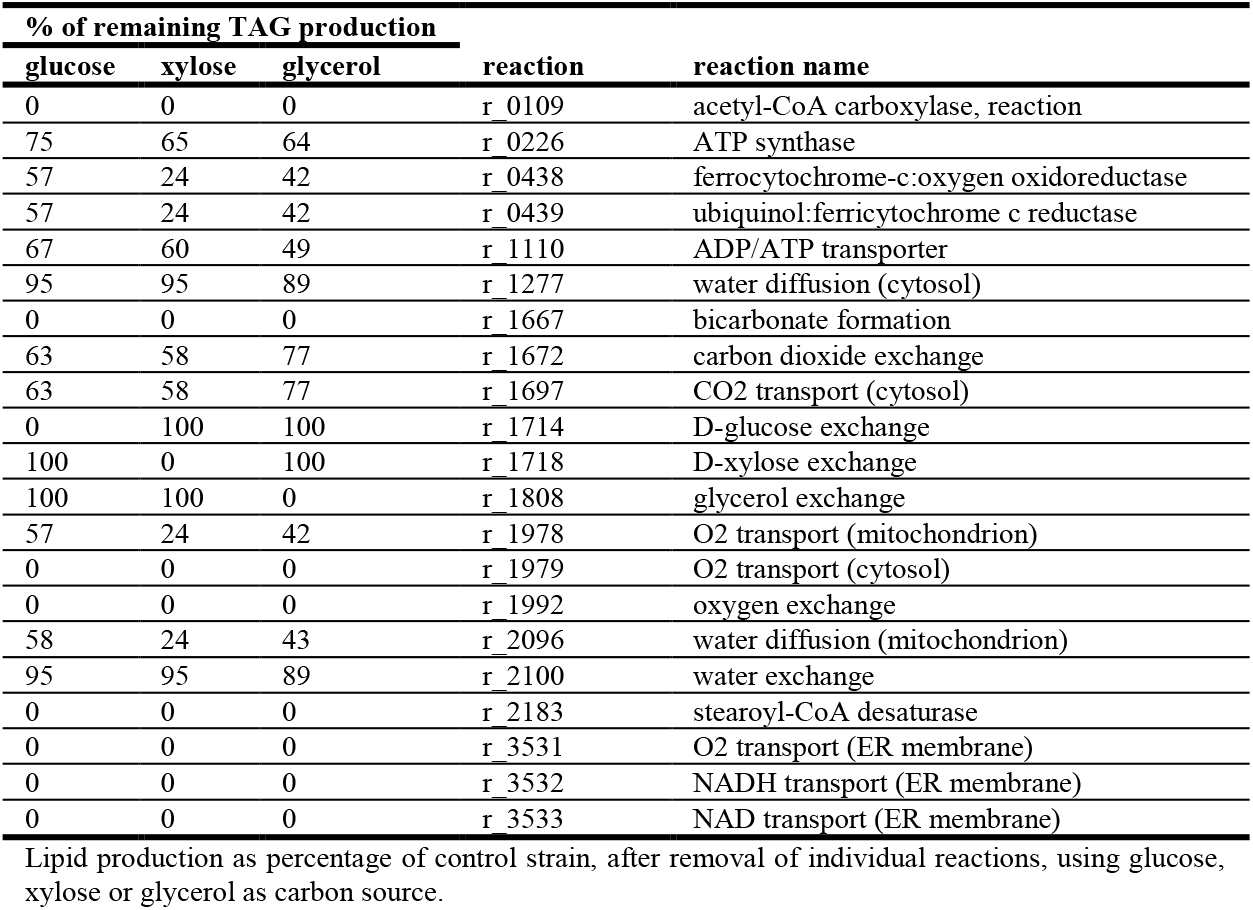
Effect on TAG production after removal of reactions important for oleaginous phenotype.

### 3.6 Prediction of potential targets for increased production of TAGs

Genetic studies of *R. toruloides* have shown that its lipid accumulation capacity can be further enhanced (reviewed in (Marella, Holkenbrink, Siewers, & Borodina, 2018)). We employed Flux Scanning based on Enforced Objective Function (FSEOF) on *rhto-GEM* to identify potential metabolic engineering targets for improved production of TAG. FSEOF is based on the principle that increased production requires a redirection of flux, from originally going towards biomass generation, to ideally going (partially) towards our product of interest. However, reactions that already carry significant flux for biomass production are accounted for in FSEOF, as these are potentially less promising targets for overexpression. As there is interest in the use of hydrolysed plant biomass as feedstock, we evaluated potential targets for both glucose and xylose as carbon source, in addition to the often used glycerol (**Table 2, Fig 3**).

**Fig. 3.**
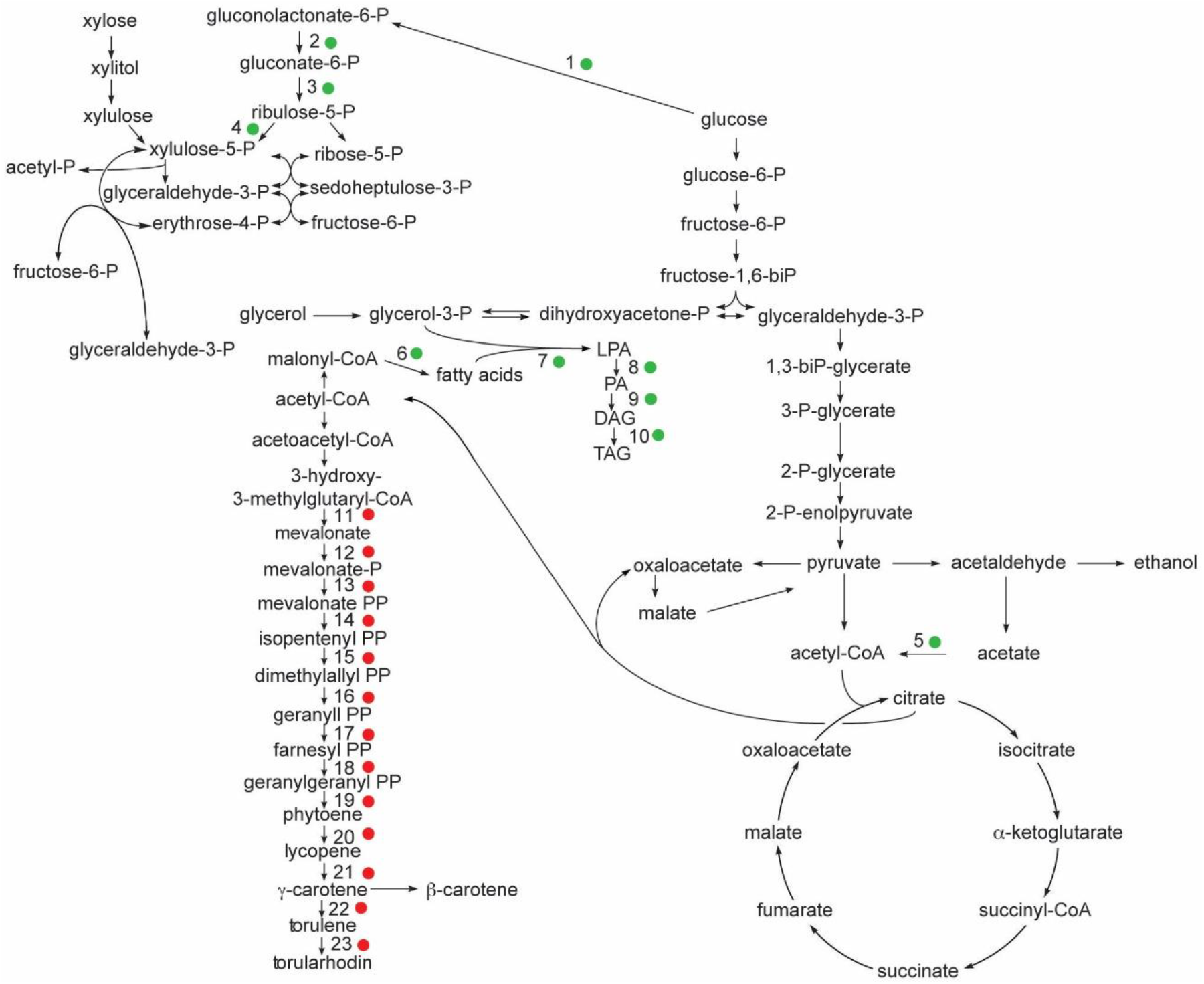
Targets predicted by FSEOF. Genetic engineering targets for improved production of TAGs (green circles) and torularhodin (red circles) in glucose-grown *R. toruloides*. The numbers refer to reactions indicated in Table 2 and 3.

**Table 2.**
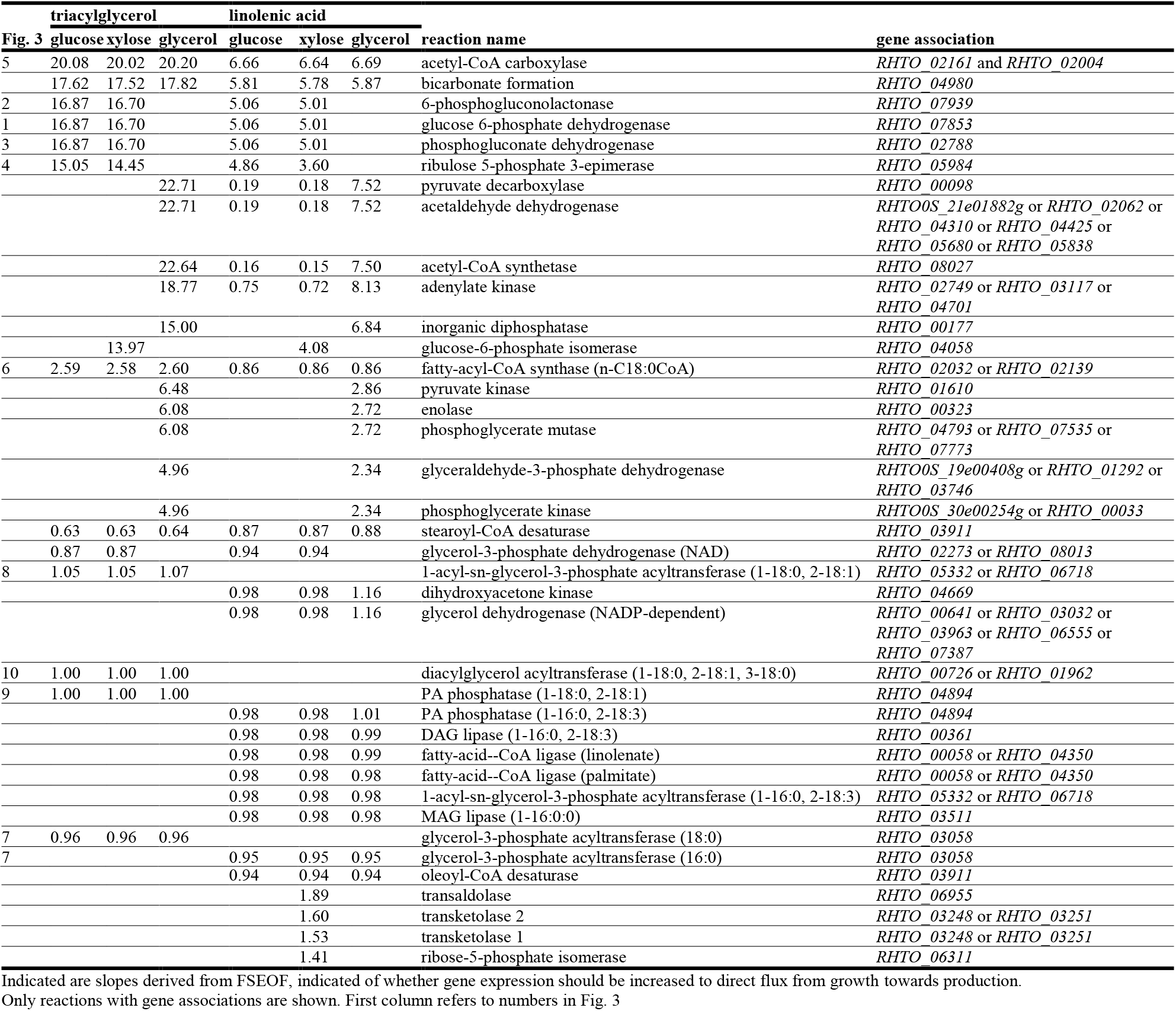
Comparison of targets predicted from FSEOF for improved TAG and linolenic production on glucose, xylose or glycerol as carbon source.

Many glycolytic genes were predicted as genetic engineering targets for improved production of TAGs, while enzymes of the pentose phosphate (PP) pathway were also identified as targets when cultivated on xylose, as anticipated. All three components of the pyruvate decarboxylase (*PDC*)-aldehyde dehydrogenase (*ALD*)-acetyl-CoA synthetase (*ACS*) pathway were predicted to be targets suitable for overexpression under cultivation on xylose, which was unexpected as this pathway is considered less energy efficient than the pyruvate dehydrogenase (*PDH*)-ATP-citrate lyase (*ACL*) pathway which is active in oleaginous yeasts. However, in the non-oleaginous fermentative yeast *S. cerevisiae* upregulation of different members of the *PDH* bypass was shown to improve TAG accumulation (Shiba, Paradise, Kirby, Ro, & Keasling, 2007), and upregulation of the *PDH* bypass also improved lipid production in *Y. lipolytica* (Xu, Qiao, Ahn, & Stephanopoulos, 2016).

More of the identified targets have previously been validated by experimental results: i.e. overexpression of native acetyl-CoA carboxylase (ACC1), diacylglycerol O-acyltransferase (*DGA1*), glycerol-3-phosphate dehydrogenase (*GUT2*) genes have all been shown to increase TAG yield in *R. toruloides* (S. Zhang et al., 2016), while increased expression of fatty-acyl-CoA synthase (*FAS1* and *FAS2*) was beneficial in *S. cerevisiae* (Runguphan & Keasling, 2014). Similarly, the prediction that upregulation of stearoyl-CoA desaturase (*OLE1*) agreed with a published report that overexpression of this enzyme increases lipid production (S. Zhang et al., 2016). This relates to fatty acid composition of TAGs in *R. toruloides*, mostly containing oleic acid at sn-2 position, as prevailing TAGs (~15% each) are POO (C16:0, C18:1, C18:1), POP (C16:0, C18:1, C16:0), POS (C16:0, C18:1, C18:0) and minor TAGs (10-5% each) are PLO (C16:0, C18:2, C18:1), PLP (C16:0, C18:2, C16:0), PLS (C16:0, C18:2, C18:0) (Wei et al., 2017). Desaturation of palmitic and stearic acid, which may inhibit acetyl-CoA synthase, has been shown to produce higher lipid yields in *Y. lipolytica* (Qiao, Wasylenko, Zhou, Xu, & Stephanopoulos, 2017).

Gene targets that have previously been shown to enhance TAG accumulation in *R. toruloides* but were not predicted in the current analysis include non-metabolic genes such as those involved in organelle morphogenesis, which are currently beyond the capacity of analysis of a purely metabolic model. This include genes such as *RHTO_05627*, which encodes the lipid droplet-associated protein Ldp1 (Zhu et al., 2015) whose expression has been shown to improve lipid production.

### 3.7 Prediction of potential targets for increased production of linolenic acid

Oleaginous yeasts are a potential source of essential fatty acids such as linolenic and linoleic acid, while linolenic ω-3 fatty acids provide health benefits in nutrition and serve as precursors for synthesis of docosahexaenoic acid (DHA) and eicosapentaenoic acid (EPA) in humans (Innis, 2014). Natural strains of *R. toruloides* contain linolenic acid around 3% of total fatty acids and the level of fatty acids saturation can change in response to temperature (Suutari, Liukkonen, & Laakso, 1990). Genetic engineering targets for improved production of polyunsatured fatty acids (PUFAs) in microorganisms have been reviewed previously (Gong et al., 2014). We performed FSEOF analysis specifically for linolenic acid production, and as expected most genes identified as targets were also identified for improved production of TAGs, in addition to oleoyl-CoA and linoleoyl-CoA desaturases (**Table 2**). This is furthermore supported by published reports, e.g., overexpression of native Δ9 desaturase (Tsai et al., 2019; S. Zhang et al., 2016) was shown to result in increase of linolenic acid production in *R. toruloides*.

In the yeast cell PUFAs may occur as constituent of phospholipids, sulfolipids, acylglycerols or glycolipids (Jacob, 1992). Fungal Δ6-desaturases were shown to have preference for acyl groups esterified at the sn-2 position of phosphatidylcholine (PC) over acyl-CoAs. This PC-linked Δ6-desaturation pathway creates limiting step in synthesis of ω-3 fatty acids. Acyl-PCs may be channelled towards TAG synthesis via acyltransferase reaction.

Understanding of substrate specificity of desaturases and acyltransferases could contribute to regulation of PC, acyl-CoAs, TAGs pools and more accurate determination of fluxes that should be enhanced for improved production of linolenic acid. Overexpression of heterologous desaturases (with desired substrate specificity) from organisms naturally overproducing ω-3 fatty acids can be a promising approach as compared to overexpression of native desaturases (Wang, 2013). In *Y. lipolytica* overexpression of heterologous desaturases resulted in enhanced production of ω-3 fatty acids with EPA as final product at the highest content among known EPA sources (Xie, Jackson, & Zhu, 2015; Xue et al., 2013).

### 3.8 Prediction of potential targets for improved production of carotenoids

Carotenoids are terpenoid pigments that are widely used as natural colorants in food industry (Mata-Gómez, Montañez, Méndez-Zavala, & Aguilar, 2014). Several genera of basidiomycete yeast are natural producers of carotenoids including *Sporobolomyces*, *Sporidiobolus*, *Rhodotorula* and *Xanthophyllomyces/Phaffia* (Buzzini et al., 2007). Many of these yeasts reside in the phylloplane and their ability to produce carotenoids is thought to serve as protection against solar radiation.

Carotenoid production in yeast has been shown to depend on cultivation conditions (Dias, Silva, Freitas, Reis, & da Silva, 2016; Singh et al., 2016) with yields of 0.28 mg/g having been achieved in fed-batch cultivation of *R. toruloides* (Dias, Sousa, Caldeira, Reis, & Lopes da Silva, 2015). *R. toruloides* produces a number of carotenoids including torularhodin, torulene, γ-carotene and β-carotene. Torulene is the major carotenoid, comprising 50 % (w/w) of pigment produced, followed by torularhodin and γ-carotene, which account for 20 % each. Torulene, which contains 13 double bonds per molecule, has shown to display higher antioxidant activity than β-carotene, which contains 11 double bonds per molecule (Sakaki, Nochide, Komemushi, & Miki, 2002).

We performed FSEOF analysis to predict targets for carotenoid production, with torularhodin as representative product (**Table 3, Fig 3**). Most of enzymes within the mevalonate pathway were predicted as targets for overexpression, as this pathway is responsible for the production of the prenyl pyrophosphate precursor. This indicates that a higher flux is required for carotenoids than what is obtained during biomass production. Overexpression of truncated 3-hydroxy-3-methylglutaryl-CoA reductase from *Kluyveromyces marxianus* in combination with other genes was shown to increase β-carotene production in *Rhodotorula glutinis* (Pi et al., 2018). Similarly, also all enzymes of isoprene biosynthetic pathway were predicted to correlate with improved carotenoid production in *R. toruloides*.

**Table 3.** Comparison of targets predicted from FSEOF for improved torularhodin production on glucose, xylose or glycerol as carbon source.

| torularhodin |  |  |  | reaction name | gene association |
| --- | --- | --- | --- | --- | --- |
| Fig. 3 | glucose | xylose | glycerol |  |  |
| 12 | 3.96 | 3.95 | 3.96 | mevalonate kinase (atp) | *RHTO_02122* |
| 11 | 3.95 | 3.95 | 3.95 | hydroxymethylglutaryl CoA reductase | *RHTO_04045* |
| 14 | 3.95 | 3.95 | 3.95 | mevalonate pyrophosphate decarboxylase | *RHTO_06005* |
| 13 | 3.95 | 3.95 | 3.95 | phosphomevalonate kinase | *RHTO_02073* |
|  | 3.51 | 4.11 |  | ribulose 5-phosphate 3-epimerase | *RHTO_05984* |
|  | 1.67 | 1.60 |  | 6-phosphogluconolactonase | *RHTO_07939* |
|  | 1.67 | 1.60 |  | glucose 6-phosphate dehydrogenase | *RHTO_07853* |
|  | 1.67 | 1.60 |  | phosphogluconate dehydrogenase | *RHTO_02788* |
| 18 | 1.00 | 1.00 | 1.00 | farnesyltranstransferase | *RHTO_01660* or *RHTO_02504* |
| 19, 21, 22 | 1.00 | 1.00 | 1.00 | phytoene synthase | *RHTO_04605* |
| 20, 23 | 1.00 | 1.00 | 1.00 | phytoene dehydrogenase | *RHTO_04602* |
| 16 | 0.98 | 0.98 | 0.98 | dimethylallyltranstransferase | *RHTO_01660* |
| 17 | 0.98 | 0.98 | 0.98 | geranyltranstransferase | *RHTO_01660* |
| 15 | 0.98 | 0.98 | 0.98 | isopentenyl-diphosphate D-isomerase | *RHTO_05138* |
|  |  |  | 0.04 | dihydroxyacetone kinase | *RHTO_04669* |
|  |  |  | 0.04 | glycerol dehydrogenase (NADP-dependent) | *RHTO_00641* or *RHTO_03032* or *RHTO_03963* or *RHTO_06555* or *RHTO_07387* |

The dimethylallyltranstransferase/geranyltranstransferase *(ERG20)* is involved in synthesis of farnesyl pyrophosphate (FPP) from geranyl pyrophosphate (GPP) and IPP, which is a direct metabolic precursor of carotenoids as well as ergosterol, heme A, dolichols and prenyl-adducts for prenylated proteins. A recent study has demonstrated that overexpression of the *X. dendrorhous* geranylgeranyl pyrophosphate (GGPP) synthase (encoded by the *BTS1* gene), which is immediately downstream of *ERG20*, increased carotenoid production in *R. glutinis* (Pi et al., 2018).

Carotenoid biosynthetic enzymes, such as phytoene synthase and dehydrogenase were identified as targets for increased carotenoid production, as anticipated. The gene coding for phytoene dehydrogenase (*RHTO_04602*) has been experimentally verified to be involved in carotenoid biosynthesis (Sun et al., 2017). In addition, overexpression of the *X. dendrorhous* phytoene desaturase and synthase genes was shown to increase carotenoid yields in *R. glutinis* (Pi et al., 2018).

Little genetic analysis of the endogenous enzymes for carotenoid biosynthesis in *R. toruloides* has been carried out to date. A number of mutants with improved carotenoid yields have been generated but the exact genetic changes responsible have not been reported (Bao et al., 2019; C. Zhang et al., 2016) The earlier mentioned T-DNA mutagenesis study reported decreased carotenoid production upon integration into the intron of *R. toruloides* hypothetical gene *RTHO_00032* or the exon of hypothetical gene *RTHO_07952*, which is predicted to encode a bZIP transcription factor (Lin et al., 2017). The same study also reported that T-DNA insertion into the promoter of hypothetical gene *RTHO_07650* (encoding a putative DUF1479 domain protein) increased carotenoid yields, exemplifying that regulation plays an important role in carotenoid biosynthesis, an assertion that would benefit greatly from integrative analysis of expression data with a comprehensive model of metabolism.

Collectively, the FSEOF results have demonstrated the ability of *rhto-GEM* to provide valuable predictions of targets for improved production of key products in *R. toruloides*, as many of these have experimentally been validated to increase production. This renders *rhto-GEM* as persuasive tool to aid in improving the production of less-studied high-value compounds, in addition as a framework for more detailed analysis of high-producing strains.

## 4 Conclusion

Previous studies have presented GEMs of several oleaginous fungal species, including oleaginous ascomycete *Yarrowia lipolytica* (Loira, Dulermo, Nicaud, & Sherman, 2012), zygomycetes *Mortierella alpina* (Ye et al., 2015) and *Mucor circinelloides* (Vongsangnak et al., 2016). Our study presents the first reconstruction of GEM of lipid-accumulating basidiomycete *R. toruloides*, and while we used the *S. cerevisiae* model as reference for the conserved parts of metabolism, *rhto-GEM* contains unique characteristics including ATP:citrate lyase, which is the main source of acetyl-CoA for lipid synthesis; mitochondrial beta-oxidation; a cytoplasmic malic enzyme that provides an alternative to the PP pathway for NADPH regeneration; and pathways related to poly-unsaturated fatty acids and carotenoid biosynthesis.

The model incorporates knowledge obtained from genomics and proteomics data generated for *R. toruloides* (Zhu et al., 2012) and was validated using cultivation data (Azambuja et al., 2018; Bommareddy et al., 2015; Bonturi et al., 2017; Shen et al., 2013), demonstrating good agreement with experimentally reported growth rates. Analysis of the model allowed to identify potential genetic engineering strategies for enhanced lipid production. Some of these genetic targets were found to agree with published experimental studies (Díaz et al., 2018; S. Zhang et al., 2016). As such, *rhto-GEM* emerges as a valuable tool for future analysis of oleaginous and lipid metabolism. An important feature is its distribution through a Git repository, which allows for continuous improvement and tracking of model development.

By providing all relevant scripts to replicate the reconstruction of *rhto-GEM* we have demonstrated how a new genome-scale model can conveniently be generated, a process greatly aided by RAVEN or alternative solutions such as AutoKEGGRec (Karlsen, Schulz, & Almaas, 2018). Expeditious automated reconstruction of a draft model is followed by manual curation to produce a model accurately representing the *in vivo* metabolic network. This step remains the most time-consuming involving literature study and comparison of simulations with reported experimental results. For lesser studied organisms, such as many of the oleaginous yeasts, this step has the additional challenge of limited available experimental data. Fortunately, continuous research interest in oleaginous yeasts generates new data and knowledge and this can subsequently be applied to curate and iteratively improve the existing model. As such, a genome-scale model is never *finished*, it is merely describing the current knowledge. To facilitate this, models should be versioned, and their curation tracked in an accessible development environment, while the quality of each model version should be assured by e.g. a dedicated testing suite as memote. Through such an approach emerging knowledge on *R. toruloides* metabolism can easily and reproducibly be used to push *rhto-GEM* as a comprehensive tool, which through its open nature is exemplary suited for involvement of other researchers inside the *R. toruloides* community. A move towards free distribution of model improvements at an early stage will be of strong benefit for the research community, while tracking of changes retains the ability to give credit to the responsible contributors.

## Acknowledgements

The authors acknowledge Dr. Benjamín José Sánchez (Chalmers University of Technology) for valuable discussions and Dr. Tomas Linder (Swedish University of Agricultural Sciences) for assistance with the figures.

## Supporting Information

**Supplementary Material S1. Memote report *rhto-GEM* v1.2.1**

## References

Andrade, R., Leal, R., Roseiro, J., Reis, A., & da Silva, T. L. (2012). Monitoring Rhodosporidium toruloides NCYC 921 batch fermentations growing under carbon and nitrogen limitation by flow cytometry. World Journal of Microbiology and Biotechnology, 28(3), 1175–1184. https://doi.org/10.1007/s11274-011-0920-2

Azambuja, S. P. H., Bonturi, N., Miranda, E. A., & Gombert, A. K. (2018). Physiology and lipid accumulation capacity of different Yarrowia lipolytica and Rhodosporidium toruloides strains on glycerol. BioRxiv, 1–18. https://doi.org/10.1101/278523

Bao, R., Gao, N., Lv, J., Ji, C., Liang, H., Li, S., … Lin, X. (2019). Enhancement of Torularhodin Production in Rhodosporidium toruloides by Agrobacterium tumefaciens-Mediated Transformation and Culture Condition Optimization. Journal of Agricultural and Food Chemistry, 67(4), 1156–1164. research-article. https://doi.org/10.1021/acs.jafc.8b04667

Bommareddy, R. R., Sabra, W., Maheshwari, G., & Zeng, A.-P. (2015). Metabolic network analysis and experimental study of lipid production in Rhodosporidium toruloides grown on single and mixed substrates. Microbial Cell Factories, 14(1), 36. https://doi.org/10.1186/s12934-015-0217-5

Bonturi, N., Crucello, A., Viana, A. J. C., & Miranda, E. A. (2017). Microbial oil production in sugarcane bagasse hemicellulosic hydrolysate without nutrient supplementation by a Rhodosporidium toruloides adapted strain. Process Biochemistry, 57, 16–25. https://doi.org/10.1016/j.procbio.2017.03.007

Buzzini, P., Innocenti, M., Turchetti, B., Libkind, D., van Broock, M., & Mulinacci, N. (2007). Carotenoid profiles of yeasts belonging to the genera *Rhodotorula*, *Rhodosporidium*, *Sporobolomyces*, and *Sporidiobolus*. Canadian Journal of Microbiology, 53(8), 1024–1031. https://doi.org/10.1139/W07-068

Camacho, C., Coulouris, G., Avagyan, V., Ma, N., Papadopoulos, J., Bealer, K., & Madden, T. L. (2009). BLAST+: architecture and applications. BMC Bioinformatics, 10(1), 421. https://doi.org/10.1186/1471-2105-10-421

Castañeda, M. T., Nuñez, S., Garelli, F., Voget, C., & De Battista, H. (2018). Comprehensive analysis of a metabolic model for lipid production in Rhodosporidium toruloides. Journal of Biotechnology, 280, 11–18. https://doi.org/10.1016/j.jbiotec.2018.05.010

Choi, H. S., Lee, S. Y., Kim, T. Y., & Woo, H. M. (2010). In Silico Identification of Gene Amplification Targets for Improvement of Lycopene Production. Applied and Environmental Microbiology, 76(10), 3097–3105. https://doi.org/10.1128/AEM.00115-10

Coradetti, S. T., Pinel, D., Geiselman, G. M., Ito, M., Mondo, S. J., Reilly, M. C., … Skerker, J. M. (2018). Functional genomics of lipid metabolism in the oleaginous yeast Rhodosporidium toruloides. ELife, 7, e32110. https://doi.org/10.7554/eLife.32110

Dias, C., Silva, C., Freitas, C., Reis, A., & da Silva, T. L. (2016). Effect of Medium pH on Rhodosporidium toruloides NCYC 921 Carotenoid and Lipid Production Evaluated by Flow Cytometry. Applied Biochemistry and Biotechnology, 179(5), 776–787. https://doi.org/10.1007/s12010-016-2030-y

Dias, C., Sousa, S., Caldeira, J., Reis, A., & Lopes da Silva, T. (2015). New dual-stage pH control fed-batch cultivation strategy for the improvement of lipids and carotenoids production by the red yeast Rhodosporidium toruloides NCYC 921. Bioresource Technology, 189, 309–318. https://doi.org/10.1016/j.biortech.2015.04.009

Díaz, T., Fillet, S., Campoy, S., Vázquez, R., Viña, J., Murillo, J., & Adrio, J. L. (2018). Combining evolutionary and metabolic engineering in Rhodosporidium toruloides for lipid production with non-detoxified wheat straw hydrolysates. Applied Microbiology and Biotechnology, 102(7), 3287–3300. https://doi.org/10.1007/s00253-018-8810-2

Gong, Y., Wan, X., Jiang, M., Hu, C., Hu, H., & Huang, F. (2014). Metabolic engineering of microorganisms to produce omega-3 very long-chain polyunsaturated fatty acids. Progress in Lipid Research, 56(1), 19–35. https://doi.org/10.1016/j.plipres.2014.07.001

Heirendt, L., Arreckx, S., Pfau, T., Mendoza, S. N., Richelle, A., Heinken, A., … Fleming, R. M. T. (2017). Creation and analysis of biochemical constraint-based models: the COBRA Toolbox v3.0. ArXiv, 1710.04038v2.

Innis, S. M. (2014). Omega-3 Fatty Acid Biochemistry: Perspectives from Human Nutrition. Military Medicine, 179(11S), 82–87. https://doi.org/10.7205/milmed-d-14-00147

Jacob, Z. (1992). Yeast Lipids: Extraction, Quality Analysis, and Acceptability. Critical Reviews in Biotechnology, 12(5–6), 463–491. https://doi.org/10.3109/07388559209114236

Karlsen, E., Schulz, C., & Almaas, E. (2018). Automated generation of genome-scale metabolic draft reconstructions based on KEGG. BMC Bioinformatics, 19(1), 1–11. https://doi.org/10.1186/s12859-018-2472-z

Kerkhoven, E. J., Lahtvee, P.-J., & Nielsen, J. (2015). Applications of computational modeling in metabolic engineering of yeast. FEMS Yeast Research, 15(1), 1–13. https://doi.org/10.1111/1567-1364.12199

Kerkhoven, E. J., Pomraning, K. R., Baker, S. E., & Nielsen, J. (2016). Regulation of amino-acid metabolism controls flux to lipid accumulation in Yarrowia lipolytica. NPJ Systems Biology and Applications, 2, 16005. https://doi.org/10.1038/npjsba.2016.5

Lee, N. K., Cheon, C. J., & Rhee, J.-K. (2018). Anti-Obesity Effect of Red Radish Coral Sprout Extract by Inhibited Triglyceride Accumulation in a Microbial Evaluation System and in High-Fat Diet-Induced Obese Mice. Journal of Microbiology and Biotechnology, 28(3), 397–400. https://doi.org/10.4014/jmb.1802.02005

Lieven, C., Beber, M. E., Olivier, B. G., Bergmann, F. T., Chauhan, S., Correia, K., … Jasper, J. (2018). Memote: A community driven effort towards a standardized genome-scale metabolic model test suite. BioRxiv, 1–26. https://doi.org/10.1101/350991

Lin, X., Gao, N., Liu, S., Zhang, S., Song, S., Ji, C., … Zhu, B. (2017). Characterization the carotenoid productions and profiles of three Rhodosporidium toruloides mutants from Agrobacterium tumefaciens-mediated transformation. Yeast, 34(8), 335–342. https://doi.org/10.1002/yea.3236

Loira, N., Dulermo, T., Nicaud, J.-M., & Sherman, D. J. (2012). A genome-scale metabolic model of the lipid-accumulating yeast Yarrowia lipolytica. BMC Systems Biology, 6(1), 35. https://doi.org/10.1186/1752-0509-6-35

Lu, H., Li, F., Sánchez, B. J., Zhu, Z., Li, G., Domenzain, I., … Nielsen, J. (2019). Yeast8: the consensus metabolic model of S. cerevisiae and its ecosystem of models for probing cell metabolism at a multi-scale level. Submitted.

Marella, E. R., Holkenbrink, C., Siewers, V., & Borodina, I. (2018). Engineering microbial fatty acid metabolism for biofuels and biochemicals. Current Opinion in Biotechnology, 50(Table 1), 39–46. https://doi.org/10.1016/j.copbio.2017.10.002

Mata-Gómez, L., Montañez, J., Méndez-Zavala, A., & Aguilar, C. (2014). Biotechnological production of carotenoids by yeasts: an overview. Microbial Cell Factories, 13(1), 12. https://doi.org/10.1186/1475-2859-13-12

Nguyen, L. N., Bormann, J., Le, G. T. T., Stärkel, C., Olsson, S., Nosanchuk, J. D., … Schäfer, W. (2011). Autophagy-related lipase FgATG15 of Fusarium graminearum is important for lipid turnover and plant infection. Fungal Genetics and Biology, 48(3), 217–224. https://doi.org/10.1016/j.fgb.2010.11.004

Pan, J., Hu, C., & Yu, J.-H. (2018). Lipid Biosynthesis as an Antifungal Target. Journal of Fungi (Basel, Switzerland), 4(2), 72. https://doi.org/10.3390/jof4020050

Papanikolaou, S., & Aggelis, G. (2011). Lipids of oleaginous yeasts. Part I: Biochemistry of single cell oil production. European Journal of Lipid Science and Technology, 113(8), 1031–1051. https://doi.org/10.1002/ejlt.201100014

Park, Y.-K., Nicaud, J.-M., & Ledesma-Amaro, R. (2018). The Engineering Potential of Rhodosporidium toruloides as a Workhorse for Biotechnological Applications. Trends in Biotechnology, 36(3), 304–317. https://doi.org/10.1016/j.tibtech.2017.10.013

Pi, H.-W., Li, W.-H., Lin, Y.-J., Chang, J.-J., Anandharaj, M., & Kao, Y.-Y. (2018). Engineering the oleaginous red yeast Rhodotorula glutinis for simultaneous β-carotene and cellulase production. Scientific Reports, 8(1), 2–11. https://doi.org/10.1038/s41598-018-29194-z

Prigent, S., Frioux, C., Dittami, S. M., Thiele, S., Larhlimi, A., Collet, G., … Siegel, A. (2017). Meneco, a Topology-Based Gap-Filling Tool Applicable to Degraded Genome-Wide Metabolic Networks. PLoS Computational Biology, 13(1), 1–32. https://doi.org/10.1371/journal.pcbi.1005276

Pryszcz, L. P., Huerta-Cepas, J., & Gabaldón, T. (2011). MetaPhOrs: Orthology and paralogy predictions from multiple phylogenetic evidence using a consistency-based confidence score. Nucleic Acids Research, 39(5). https://doi.org/10.1093/nar/gkq953

Qiao, K., Wasylenko, T. M., Zhou, K., Xu, P., & Stephanopoulos, G. (2017). Lipid production in Yarrowia lipolytica is maximized by engineering cytosolic redox metabolism. Nature Biotechnology, 35(2), 173–177. https://doi.org/10.1038/nbt.3763

Runguphan, W., & Keasling, J. D. (2014). Metabolic engineering of Saccharomyces cerevisiae for production of fatty acid-derived biofuels and chemicals. Metabolic Engineering, 21, 103–113. https://doi.org/10.1016/j.ymben.2013.07.003

Sakaki, H., Nochide, H., Komemushi, S., & Miki, W. (2002). Effect of active oxygen species on the productivity of torularhodin by Rhodotorula glutinis no. 21. Journal of Bioscience and Bioengineering, 93(3), 338–340. https://doi.org/10.1016/S1389-1723(02)80040-8

Sampaio, J. P. (2011). RhodosporidiumBanno (1967). The Yeasts, 3(1967), 1523–1539. https://doi.org/10.1016/B978-0-444-52149-1.00127-0

Sánchez, B. J., Li, F., Kerkhoven, E. J., & Nielsen, J. (2019). SLIMEr: probing flexibility of lipid metabolism in yeast with an improved constraint-based modeling framework. BMC Systems Biology, 13(1), 4. https://doi.org/10.1186/s12918-018-0673-8

Sánchez, B. J., Li, F., Lu, H., Kerkhoven, E. J., & Nielsen, J. (2018). SysBioChalmers/yeast-GEM: yeast 8.2.0. Zenodo. https://doi.org/10.5281/zenodo.1495483

Shen, H., Gong, Z., Yang, X., Jin, G., Bai, F., & Zhao, Z. K. (2013). Kinetics of continuous cultivation of the oleaginous yeast Rhodosporidium toruloides. Journal of Biotechnology, 168(1), 85–89. https://doi.org/10.1016/j.jbiotec.2013.08.0102

Shi, J., Feng, H., Lee, J., & Chen, W. N. (2013). Comparative proteomics profile of lipid-cumulating oleaginous yeast: An iTRAQ-coupled 2-D LC-MS/MS analysis. PLoS ONE, 8(12), e85532. https://doi.org/10.1371/journal.pone.0085532

Shiba, Y., Paradise, E. M., Kirby, J., Ro, D. K., & Keasling, J. D. (2007). Engineering of the pyruvate dehydrogenase bypass in Saccharomyces cerevisiae for high-level production of isoprenoids. Metabolic Engineering, 9(2), 160–168. https://doi.org/10.1016/j.ymben.2006.10.005

Singh, G., Jawed, A., Paul, D., Bandyopadhyay, K. K., Kumari, A., & Haque, S. (2016). Concomitant Production of Lipids and Carotenoids in Rhodosporidium toruloides under Osmotic Stress Using Response Surface Methodology. Frontiers in Microbiology, 7(October), 1–13. https://doi.org/10.3389/fmicb.2016.01686

Soccol, C. R., Dalmas Neto, C. J., Soccol, V. T., Sydney, E. B., da Costa, E. S. F., Medeiros, A. B. P., & Vandenberghe, L. P. de S. (2017). Pilot scale biodiesel production from microbial oil of Rhodosporidium toruloides DEBB 5533 using sugarcane juice: Performance in diesel engine and preliminary economic study. Bioresource Technology, 223, 259–268. https://doi.org/10.1016/j.biortech.2016.10.055

Sun, W., Yang, X., Wang, X., Lin, X., Wang, Y., Zhang, S., … Zhao, Z. K. (2017). Homologous gene targeting of a carotenoids biosynthetic gene in Rhodosporidium toruloides by Agrobacterium-mediated transformation. Biotechnology Letters, 39(7), 1001–1007. https://doi.org/10.1007/s10529-017-2324-3

Suutari, M., Liukkonen, K., & Laakso, S. (1990). Temperature adaptation in yeasts: the role of fatty acids. Journal of General Microbiology, 136(8), 1469–1474. https://doi.org/10.1099/00221287-136-8-1469

Tchakouteu, S. S., Kalantzi, O., Gardeli, C., Koutinas, A. A., Aggelis, G., & Papanikolaou, S. (2015). Lipid production by yeasts growing on biodiesel-derived crude glycerol: strain selection and impact of substrate concentration on the fermentation efficiency. Journal of Applied Microbiology, 118(4), 911–927. https://doi.org/10.1111/jam.12736

Tiukova IA, Brandenburg J, Blomqvist J, Sampels S, Mikkelsen N, Skaugen M, et al. Proteome analysis of xylose metabolism in *Rhodotorula toruloides* during lipid production. Biotechnology for Biofuels. 2019;12(1):137. https://doi.org/10.1186/s13068-019-1478-8

Tsai, Y.-Y., Ohashi, T., Wu, C.-C., Bataa, D., Misaki, R., Limtong, S., & Fujiyama, K. (2019). Delta-9 fatty acid desaturase overexpression enhanced lipid production and oleic acid content in Rhodosporidium toruloides for preferable yeast lipid production. Journal of Bioscience and Bioengineering, 127(4), 430–440. https://doi.org/10.1016/J.JBIOSC.2018.09.005

Vongsangnak, W., Klanchui, A., Tawornsamretkit, I., Tatiyaborwornchai, W., Laoteng, K., & Meechai, A. (2016). Genome-scale metabolic modeling of Mucor circinelloides and comparative analysis with other oleaginous species. Gene, 583(2), 121–129. https://doi.org/10.1016/j.gene.2016.02.028

Vorapreeda, T., Thammarongtham, C., Cheevadhanarak, S., & Laoteng, K. (2012). Alternative routes of acetyl-CoA synthesis identified by comparative genomic analysis: involvement in the lipid production of oleaginous yeast and fungi. Microbiology, 158(1), 217–228. https://doi.org/10.1099/mic.0.051946-0

Wang, C., & St. Leger, R. J. (2007). The Metarhizium anisopliae perilipin homolog MPL1 regulates lipid metabolism, appressorial turgor pressure, and virulence. Journal of Biological Chemistry, 282(29), 21110–21115. https://doi.org/10.1074/jbc.M609592200

Wang, H., Marcišauskas, S., Sánchez, B. J., Domenzain, I., Hermansson, D., Agren, R., … Kerkhoven, E. J. (2018). RAVEN 2.0: A versatile toolbox for metabolic network reconstruction and a case study on Streptomyces coelicolor. PLOS Computational Biology, 14(10), e1006541. https://doi.org/10.1371/journal.pcbi.1006541

Wei, Y., Siewers, V., & Nielsen, J. (2017). Cocoa butter-like lipid production ability of non-oleaginous and oleaginous yeasts under nitrogen-limited culture conditions. Applied Microbiology and Biotechnology, 101(9), 3577–3585. https://doi.org/10.1007/s00253-017-8126-7

Wirth, F., & Goldani, L. Z. (2012). Epidemiology of rhodotorula: An emerging pathogen. Interdisciplinary Perspectives on Infectious Diseases, 2012. https://doi.org/10.1155/2012/465717

Wu, S., Hu, C., Jin, G., Zhao, X., & Zhao, Z. K. (2010). Phosphate-limitation mediated lipid production by Rhodosporidium toruloides. Bioresource Technology, 101(15), 6124–6129. https://doi.org/10.1016/j.biortech.2010.02.111

Wu, S., Zhao, X., Shen, H., Wang, Q., & Zhao, Z. K. (2011). Microbial lipid production by Rhodosporidium toruloides under sulfate-limited conditions. Bioresource Technology, 102(2), 1803–1807. https://doi.org/10.1016/j.biortech.2010.09.033

Xie, D., Jackson, E. N., & Zhu, Q. (2015). Sustainable source of omega-3 eicosapentaenoic acid from metabolically engineered Yarrowia lipolytica: from fundamental research to commercial production. Applied Microbiology and Biotechnology, 99(4), 1599–1610. https://doi.org/10.1007/s00253-014-6318-y

Xu, P., Qiao, K., Ahn, W. S., & Stephanopoulos, G. (2016). Engineering Yarrowia lipolytica as a platform for synthesis of drop-in transportation fuels and oleochemicals. Proceedings of the National Academy of Sciences, 113(39), 10848–10853. https://doi.org/10.1073/pnas.1607295113

Xue, Z., Sharpe, P. L., Hong, S. P., Yadav, N. S., Xie, D., Short, D. R., … Zhu, Q. (2013). Production of omega-3 eicosapentaenoic acid by metabolic engineering of Yarrowia lipolytica. Nature Biotechnology, 31(8), 734–740. https://doi.org/10.1038/nbt.2622

Ye, C., Xu, N., Chen, H., Chen, Y. Q., Chen, W., & Liu, L. (2015). Reconstruction and analysis of a genome-scale metabolic model of the oleaginous fungus Mortierella alpina. BMC Systems Biology, 9, 1. https://doi.org/10.1186/s12918-014-0137-8

Zhang, C., Shen, H., Zhang, X., Yu, X., Wang, H., Xiao, S., … Zhao, Z. K. (2016). Combined mutagenesis of Rhodosporidium toruloides for improved production of carotenoids and lipids. Biotechnology Letters, 38(10), 1733–1738. https://doi.org/10.1007/s10529-016-2148-6

Zhang, S., Skerker, J. M., Rutter, C. D., Maurer, M. J., Arkin, A. P., & Rao, C. V. (2016). Engineering Rhodosporidium toruloides for increased lipid production. Biotechnology and Bioengineering, 113(5), 1056–1066. https://doi.org/10.1002/bit.25864

Zhu, Z., Ding, Y., Gong, Z., Yang, L., Zhang, S., Zhang, C., … Zhaoa, Z. K. (2015). Dynamics of the lipid droplet proteome of the oleaginous yeast Rhodosporidium toruloides. Eukaryotic Cell, 14(3), 252–264. https://doi.org/10.1128/EC.00141-14

Zhu, Z., Zhang, S., Liu, H., Shen, H., Lin, X., Yang, F., … Zhao, Z. K. (2012). A multi-omic map of the lipid-producing yeast Rhodosporidium toruloides. Nature Communications, 3, 1112. https://doi.org/10.1038/ncomms2112

